# Sexual reproduction contributes to the evolution of resistance breaking isolates of the spinach pathogen *Peronospora effusa*

**DOI:** 10.1101/2021.11.05.467349

**Authors:** Petros Skiadas, Joël Klein, Thomas Quiroz Monnens, Joyce Elberse, Ronnie de Jonge, Guido Van den Ackerveken, Michael F. Seidl

## Abstract

*Peronospora effusa* causes downy mildew, the economically most important disease of cultivated spinach worldwide. To date, 19 *P. effusa* races have been denominated based on their capacity to break spinach resistances, but their genetic diversity and the evolutionary processes that contribute to race emergence are unknown. Here, we performed the first systematic analysis of *P. effusa* races showing that those emerge by both asexual and sexual reproduction. Specifically, we studied the diversity of 26 *P. effusa* isolates from 16 denominated races based on mitochondrial and nuclear comparative genomics. Mitochondrial genomes based on long-read sequencing coupled with diversity assessment based on short-read sequencing uncovered two mitochondrial haplogroups, each with distinct genome organization. Nuclear genome-wide comparisons of the 26 isolates revealed that ten isolates from six races could clearly be divided into three asexually evolving groups, in concordance with their mitochondrial phylogeny. The remaining isolates showed signals of reticulated evolution and discordance between nuclear and mitochondrial phylogenies, suggesting that these evolved through sexual reproduction. Increased understanding of this pathogen’s reproductive modes will provide the framework for future studies into the molecular mechanisms underlying race emergence and into the *P. effusa*-spinach interaction, thus assisting in sustainable production of spinach through knowledge-driven resistance breeding.

**Significance statement:** Many microbial plant pathogens depend on the successful colonization of their hosts to complete their life cycle, thereby damaging food crops worldwide. The most effective way of disease control is to deploy genetic disease resistances. However, the extensive use of resistant crop varieties exerts strong selective pressure on microbial plant pathogens to adapt in order to escape resistance. Through yet unknown mechanisms, the spinach pathogen *Peronospora effusa* can rapidly break the resistance of newly introduced varieties, often within a single growing season. Thus, there is an urgent need to better understand the mechanisms driving adaptation in *P. effusa*. This information will lead the way to knowledge-driven resistance breeding. Here, we capture for the first time the genetic variation of 26 *P. effusa*, 16 of which can break a different combination of host resistances. We demonstrate that *P. effusa* isolates evolve by both asexual and sexual reproduction, and thereby provide the framework to study the molecular mechanisms of the interactions between *P. effusa* and spinach.

## Introduction

Many microbial plant pathogens need to successfully colonize their hosts to complete their life cycle. Colonization leads to extensive damage and disease on the infected host plants (McMullan *et al*., 2015). To mitigate these effects, plants have evolved defence mechanisms to resist pathogens, while in turn pathogens evolve to overcome the host immune system to successfully establish an infection (Cook *et al*., 2015). In natural populations, host and pathogen populations are slowly coevolving and maintain diversity of resistance and virulence alleles (Barrett *et al*., 2009; Möller and Stukenbrock, 2017). In the last decades, modern agricultural practices have introduced resistant host cultivars that are often deployed in monocultures (Miller *et al*., 2020). This practice exerts strong evolutionary pressure on pathogen populations to overcome the introduced host resistance, and consequently pathogens are engaging in rapid evolutionary arms races (Möller and Stukenbrock, 2017; Mohd-Assaad *et al*., 2019).

The oomycete downy mildew *Peronospora effusa* (*Pe*) is an obligate biotrophic pathogen of spinach, and economically the most important disease of cultivated spinach worldwide (Lyon *et al*., 2016; Ribera *et al*., 2020). Severe downy mildew infection can rapidly destroy entire spinach fields, and even minor disease symptoms may require removing infected leaves before harvest, thus significantly decreasing profitability (Correll *et al*., 2011). In conventional spinach production, downy mildew is managed by the deployment of resistant cultivars and/or fungicide applications for short-term control (Koike *et al*., 1992). However, spinach production nowadays is shifting to organic practices, and consequently development and deployment of resistant cultivars are the most practical management tool (Kandel *et al*., 2020). Modern farming practices of high-density, year-round spinach production are an ideal environment for downy mildew, providing ‘green bridges’ that sustain pathogen survival (Lyon *et al*., 2016). Due to the added pressure from newly developed resistant spinach cultivars, *P. effusa* evolves rapidly and repeatedly overcomes host resistance, resulting in new epidemics and severe economic losses (Ribera *et al*., 2020).

Newly identified *P. effusa* isolates are assigned to different races based on their ability to infect a defined set of differential spinach lines developed by the International Seed Federation (Correll *et al*., 2015). These spinach differentials contain one or more resistance (*R*) loci that are known to effectively contain *P. effusa* races (Feng *et al*., 2018). New races are determined by the International Working Group on *Peronospora*, which annually evaluates newly identified *P. effusa* isolates for their capacity to overcome previously known host resistances. Downy mildew of spinach was first identified and reported in Britain in 1824 (Ribera *et al*., 2020). More than a century later, race 2 was described in 1958. Before 1990, only three races of the pathogen were known, and the disease could be well controlled (Koike *et al*., 1992). Likely driven by the extensive deployment of resistance loci (*R*-loci) in commercial spinach varieties, after 1990 the number of identified races increased tremendously, and 16 additional races have been discovered within the last 20 years (Supplementary Figure 1) (Lyon *et al*., 2016; Feng *et al*., 2018; Klein *et al*., 2020).

Like many oomycetes, *P. effusa* is a heterozygous diploid organism. Its genome sequence is relatively small compared with other oomycetes (58.6 Mb compared with for instance *Phytophthora infestans* which is 240 Mb (Haas *et al*., 2009)), and it is organised in 17 chromosomes with 53.7% repeat content (Fletcher *et al*., 2021). *P. effusa* can reproduce both asexually and sexually. During the spinach growing season, *P. effusa* reproduces asexually on infected plant tissues, releasing millions of asexual spores (Kandel *et al*., 2020). These spores are produced on leaves and other infected parts of the plant on sporangiophores that are visible with the naked eye. By the end of the growing season, *P. effusa* will also reproduce sexually, which happens when two *P. effusa* isolates of the opposite mating type (P1 and P2) co-occur in the same infected spinach tissue. Sexual reproduction results in new combinations of parental chromosomes, and this process has been thought to be a powerful driver of the emergence of new *P. effusa* races (Feng *et al*., 2018). However, *P. effusa* field isolates from various locations in the USA displayed limited genotypic diversity suggesting that these isolates are primarily the result of asexual reproduction. Nevertheless, genotypic diversity in historical isolates is likely influenced by sexual recombination (Lyon *et al*., 2016). The complete genome assembly of the *P. effusa* isolate UA202013 has been compared with closely related isolates from races 12, 13, and 14 with limited variation being observed (Fletcher *et al*., 2021). Consequently, we still know surprisingly little about genome-wide diversity between most of *P. effusa* isolates and races, as well as evolutionary processes that contribute to the emergence of new *P. effusa* races.

Mitochondrial genomes are ideal markers to study oomycete taxonomy and evolution as the mitochondrial genome has a five to ten times higher evolutionary rate compared to the nuclear genome (Choi *et al*., 2011; Yuan *et al*., 2017; Bourret *et al*., 2018). Previously, the mitochondrial genes *cox2* and *nad1* have been used to shed light on the relationship between different *P. effusa* isolates (Choi *et al*., 2011), which uncovered two distinct mitochondrial haplotype groups that could be linked to their geographical origin (Group I for Asia and Oceania and Group II for American, European, and Japan) (Choi *et al*., 2011). However, the mitochondrial phylogeny of denominated *P. effusa* races is currently unknown. Here, by using long-read sequencing data, we generated mitochondrial assemblies for four *P. effusa* isolates, including *Pe1* the first officially denominated *P. effusa* race which was isolated in the United States (Lyon *et al*., 2016). These four isolates differ in their mitochondrial genome structure, which correlates with their separation into two haplogroups. Combined with whole-genome short-read sequencing data, we developed both mitochondrial and nuclear phylogenies of 26 *P. effusa* isolates. Comparisons of both phylogenies revealed discordance and signatures of shared genetic material, which suggest that pervasive sexual recombination contributed to the evolution of resistance-breaking *P. effusa* isolates.

## Results

### *De novo P. effusa* mitochondrial genome assembly

To study the mitochondrial phylogeny of *P. effusa* isolates, we first sought to reconstruct the complete mitochondrial genome of *Pe1*, the first denominated *P. effusa* race. We performed whole-genome long-read sequencing using Oxford Nanopore sequencing technology on total genomic DNA obtained from *Pe1* spores (Supplementary Table 1B). To facilitate the assembly of the *Pe1* mitochondrial genome sequence, we recovered 19,627 reads based on sequence similarity searches to a collection of 26 publicly available oomycete mitochondrial genome sequences (Supplementary Table 1B; Supplementary Table 2). The sequencing reads were error-corrected and assembled using Canu, yielding one linear contig of 49,964 bp long. We manually curated this assembly to reconstruct a circular contig, which we further corrected for sequencing errors with high-quality Illumina short reads (Supplementary Table 1B). The resulting complete mitochondrial genome sequence of *Pe1* is a 41,316 bp circular molecule with 22.8% GC content, which is similar to mitochondrial genome assemblies previously reported for other oomycetes (Yuan *et al*., 2017; Fletcher *et al*., 2018). Based on sequence homology searches, we identified 42 protein-coding genes encoding 19 respiratory chain proteins, 16 ribosomal proteins, two protein transport proteins, and five hypothetical proteins also present in other oomycetes; of these, 12 were annotated in the negative strand and 30 in the positive strand (Figure 1, Supplementary Table 3). We observed that the majority of genes are conserved across the 26 selected mitochondrial reference sequences. Notably, we also identified three predicted genes (open reading frames; ORFs) without homologs in any of the references including *R13* and *R14*, which is comparable to the five ORFs without homology previously identified in *P. effusa* isolates *R13* and *R14* (Fletcher *et al*., 2018) and to the six to twelve genus-specific ORFs found in different *Phytophthora* species (Yuan *et al*., 2017; Cai and Scofield, 2020).

**Figure 1.**
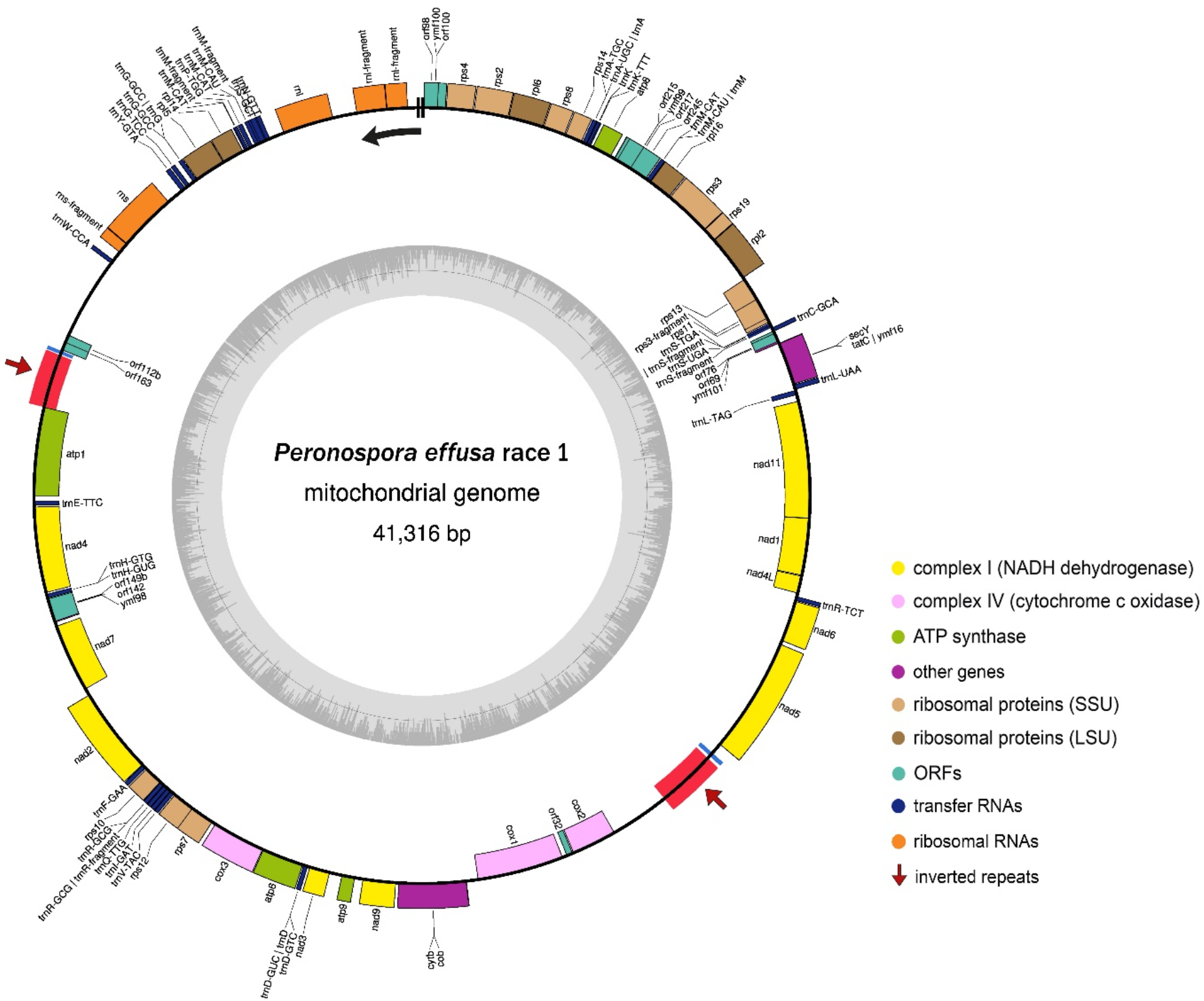
Circular representation of the mitochondrial genome assembly of *Peronospora effusa* race 1 (*Pe1*). Protein-coding genes, tRNA, rRNA, and other open-reading frames (ORFs) are shown along the outer ring (positive strand is the outside of the ring and the negative strand is the inside). The two inverted repeats are highlighted by red boxes and with arrows. The inner ring depicts the GC content. The start and end of the linear representations (Figure 2B, Figure 3B) of the circular genome assembly is indicated with two black lines at the start of the black arrow, which indicates the direction.

Mitochondrial genomes are known to contain inverted repeats (Achaz et al., 2003; Voineagu et al., 2008), and thus we used blastn to self-align the genome and identified two 100% identical 832 bp-long inverted repeats in *Pe1* (Figure 1). Inverted repeats are challenging to resolve correctly in genome assemblies when sequencing reads are not long enough to span the repetitive sequence (Wang *et al*., 2018). To corroborate the genome organization of the *Pe1* mitochondrial assembly, we mapped both short- and long-read sequencing data of *Pe1* to the assembly, which did not reveal any discordant alignments (Supplementary Figure 2), suggesting that the overall structure of the *Pe1* assembly is correct.

### Structural rearrangement in the *Peronospora effusa race 1* mitochondrial genome

Structural rearrangements in mitochondrial genomes of oomycetes are common (Yuan *et al*., 2017). We sought to investigate the mitochondrial genome structures in other Peronosporaceae and compare it to *Pe1*. To this end, we used the publicly annotated mitochondrial genome assemblies of 15 closely related species, which belong to the Peronosporaceae family, together with the *Pe1* and *R13* assembly. We first reconstructed the phylogenetic relationship between the 17 Peronosporaceae based on the mitochondrial genes *cox2* and *nad1*, which are two highly conserved genes that have been previously used as taxonomic markers to study the relationships of oomycetes including *P. effusa* (Choi *et al*., 2011). The phylogenetic tree based on concatenated *cox2* and *nad1* alignments recovers the known relationship between Peronosporaceae by clearly separating the different genera (Figure 2A).

**Figure 2.**
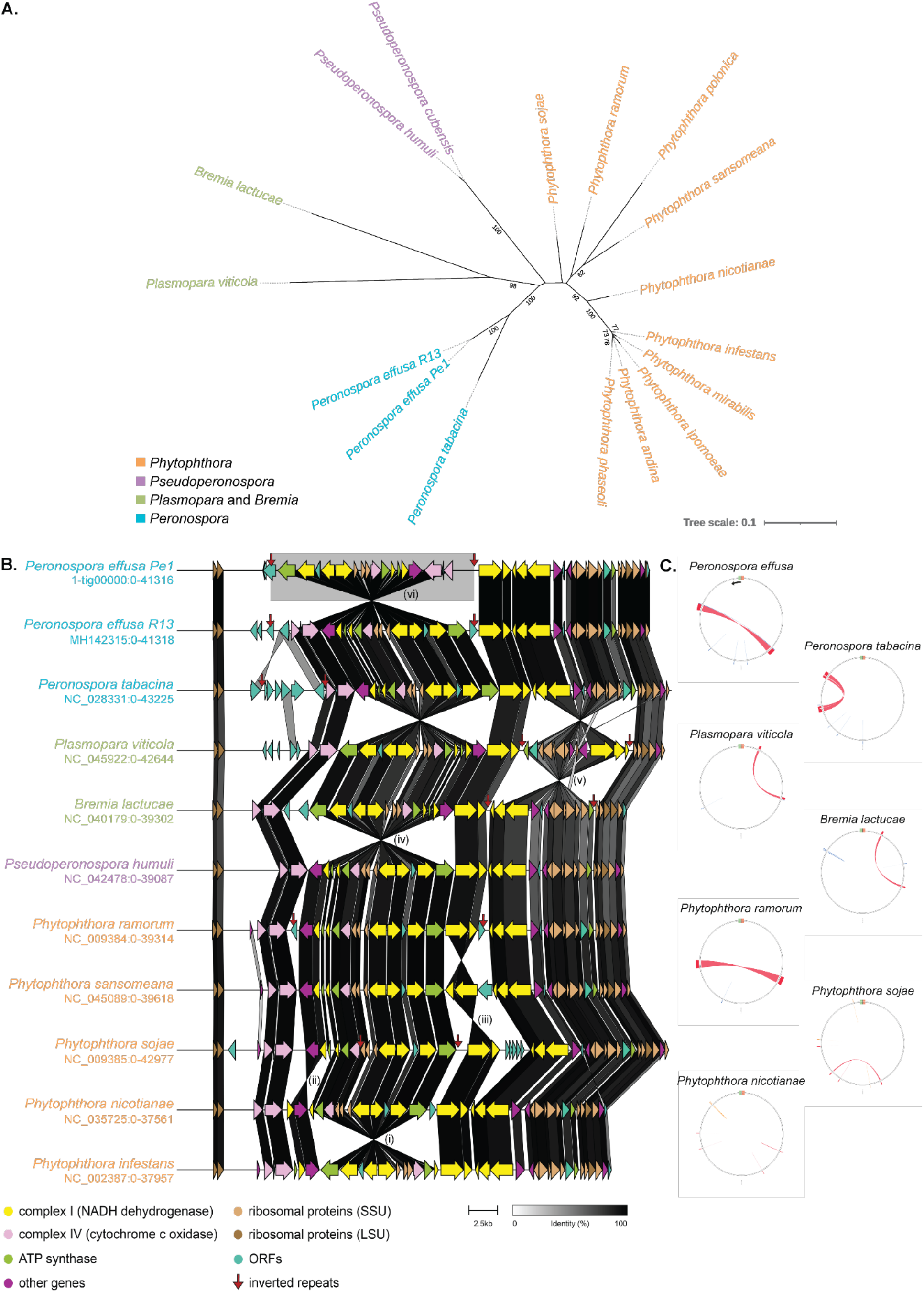
Mitochondrial genome assemblies of ten Peronosporaceae species reveal structural rearrangements that are often flanked by inverted repeats. **A)** Unrooted maximum-likelihood phylogeny of 17 different Peronosporaceae mitochondria (16 species) based on the sequences of the mitochondrial *cox2* and *nad1* genes. **B)** Linear representation of mitochondrial gene order is shown for ten Peronosporaceae, representative of the 16 species in the phylogenetic tree. Genes are coloured based on their functions and they are connected by ribbons shaded based on the percentage of protein identity (from white to black). The relative positions of the inverted repeats are shown by red arrows and the structural rearrangements are numbered (*i*-*vi*). **C)** Comparison of the inverted repeats in Peronosporaceae. The ribbons display the alignment scores, reflecting the sequence similarity (percentage): red (>75%), orange (50-75%), and blue (25%). The respective start-end of the linear representation of the circular genomes is indicated with the green/orange stripes, with the arrow indicating the direction.

Guided by their phylogenetic relationship, we reconstructed gene-based alignments between these 17 mitochondrial genomes. We observed six independent large-scale structural rearrangements across the 17 Peronosporaceae mitochondrial genomes, with *Pe1* having a unique structure that is different from *R13*; *R13* is collinear with *R14* and the publicly available assembly of the closest relative, the tobacco pathogen *Peronospora tabacina* (Fletcher *et al*., 2018). All the observed rearrangements can be summarized by displaying the differences in just ten species (Figure 2B); three of the observed structural rearrangements (*i, ii*, and *iv*) are conserved across multiple species, while three are unique to the species in which they were identified (*iii, v*, and *vi*). Two rearrangements are small (2.5 Kb) and involve only two genes (*ii, iii*), while the remaining four are larger, 8-16 Kb, and involve 11-15 genes. We found no evidence of the same region being rearranged more than once, although the regions of the rearrangements *iv* and *vi* fully overlap with those of *i* and *ii*. Thus, our data suggest that structural rearrangements are common in Peronosporaceae mitochondrial genomes, yet the organisation of mitochondrial genomes is typically conserved in closely related species. Like in *P. effusa*, we observed inverted repeats in many Peronosporaceae mitochondrial genomes. Their localization and size vary considerably between species, and they are often absent in *Phytophthora* spp. (Figure 2C), implying that they can be gained or lost multiple times independently during evolution. Notably, inverted repeats directly flank two of the unique rearrangements (*v* and *vi*), suggesting that these structures recombine to result in inversions in Peronosporaceae (Yuan *et al*., 2017).

### Mitochondrial genome phylogeny separates *Peronospora effusa* into two distinct groups

Phylogenetic analysis of the global *P. effusa* population based on the mitochondrial genes *cox2* and *nad1* has previously uncovered two distinct mitochondrial haplogroups that can be linked to the geographic origin of the individual isolates (Choi *et al*., 2011). To study the phylogenetic relationship between the denominated *P. effusa* races, we obtained spore material for 24 *P. effusa* isolates. The pathogenicity of the isolates is determined based on the current standard set of international spinach differentials (Supplementary Figure 1). To discover the evolutionary relationship of the 24 *P. effusa* isolates, we generated on average 58.8 million Illumina paired-end sequencing reads, which were subsequently mapped to the *Pe1* mitochondrial assembly to determine sequence variants (Supplementary Table 1A).

We initially focused on the mitochondrial specific genes *cox2* and *nad1* as this enabled us to study our 24 isolates together with the previously analysed 33 isolates from the global population (Choi *et al*., 2011) as well as *R13* and *R14* (Fletcher *et al*., 2018). The alignment as well as phylogenetic analyses of these genes reveal that *P. effusa* isolates can be divided into two distinct mitochondrial haplogroups (Supplementary Figure 3), thereby corroborating previous results by Choi and colleagues (Choi *et al*., 2011). Based on Choi and colleagues, European and North American isolates belong to Haplogroup II, while Asian and Australian isolates belong to Haplogroup I (Choi *et al*., 2011). However, we observed that multiple isolates denominated to Haplogroup I were isolated in the USA (*Pe5, Pe4, Pe16, US-13a*), and thus we could not establish the same correlation between the mitochondrial haplogroups and the geographical location of the *P. effusa* isolates.

To further support the separation into two haplogroups, we performed phylogenetic analyses based on the whole mitochondrial genomes of the 26 isolates (16 denominated races, eight pathotypes, as well as *R13* and *R14*). We could identify 85 variant sites (76 SNPs, 9 indels) across the mitochondrial genome in the 26 isolates based on the comparison to the *Pe1* reference mitochondrial assembly (Supplementary Table 1A). Of those, 48 variants were identified in protein-coding regions (two in ORFs that lack similarity to known protein-coding genes). Based on these 85 variants, we constructed a maximum-likelihood phylogeny, which similar to *cox2* and *nad1* supports a separation into two distinct haplogroups (separated by 46 of the 85 variants, and 100% bootstrap support); isolates within Haplogroup I differ by 21 variant sites and within Haplogroup II by 18 variant sites (Figure 3A).

**Figure 3.**
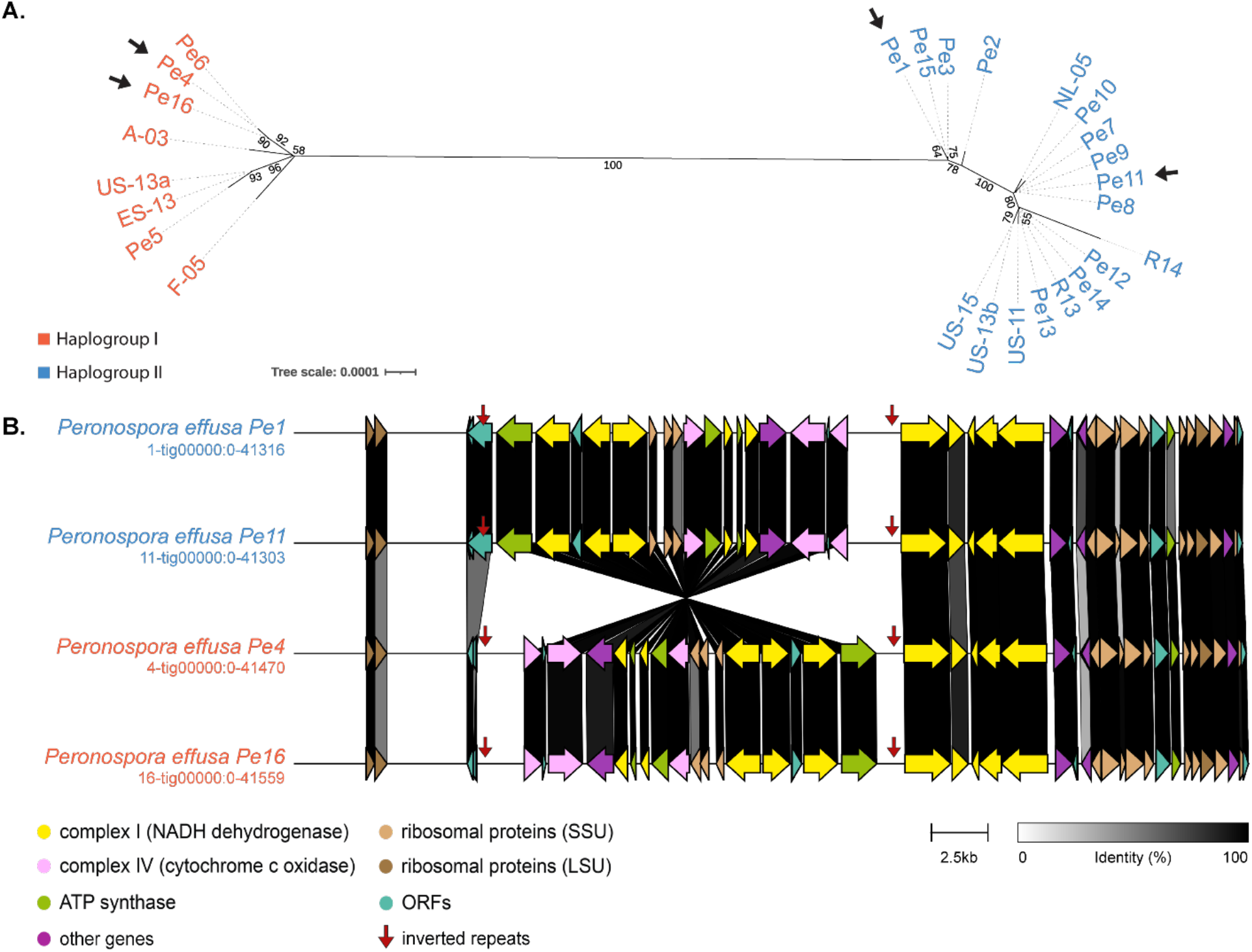
Comparison of mitochondrial genomes between *Peronospora effusa (Pe)* isolates reveals two distinct groups. **A)** Unrooted maximum-likelihood phylogeny based on the complete mitochondrial genome sequences of 26 *P. effusa* isolates clearly reveals two haplogroups (I and II). The black arrows point to the four *P. effusa* isolates, which were selected for Nanopore long-read sequencing, mitochondrial genome assembly, and gene annotation (*Pe1, Pe4, Pe6*, and *Pe11*). **B)** Mitochondrial gene orders of the selected four *P. effusa* isolates. Genes are coloured based on their functions and they are connected by ribbons coloured based on the percentage of protein identity (from white to black). The relative positions of the inverted repeats are shown by the red arrows.

Based on the discovery of the novel structure of the *Pe1* mitochondrial genome (Figure 2B), we sought to further investigate the *P. effusa* mitochondrial genome organization in the two haplogroups. To this end, we selected three additional *P. effusa* isolates for Nanopore long-read sequencing: *Pe4* and *Pe6* from Haplogroup I and *Pe11* from Haplogroup II (Supplementary Table 1B). We recovered ∽153,943 reads with an average length of 6.8 Kb based on sequence similarity searches to a collection of 26 publicly available oomycete mitochondrial genome sequences (Supplementary Table 2). We assembled the long reads with Canu and manually corrected the assemblies into circular genome sequences of 41.4 Kb on average (Supplementary Figure 4). Like in *Pe1*, these assemblies encode 43 protein-coding genes and two 100% identical, 832 bp long inverted repeats. Despite these similarities, the gene-based alignment of the four *P. effusa* mitochondrial genomes revealed a distinct genome structure for each haplogroup that is characterized by an inversion at the inverted repeats (Figure 3B). An ORF predicted on one edge of this structural rearrangement in *Pe1* and *Pe11* is missing from *Pe4* and *Pe6*. Notably, the same 16 Kb long structural rearrangement that we previously described between *Pe1* and *P. tabacina* (Figure 2B) also differentiates the two *P. effusa* haplogroups, suggesting that the structural rearrangement was introduced only in the *P. effusa* Haplogroup II.

### Nuclear and mitochondrial genome phylogenies of *Peronospora effusa* isolates are discordant

*P. effusa* reproduces both sexually and asexually (Kandel *et al*., 2020), and since the mitochondria are typically inherited from only one parental strain, mitochondrial variation alone cannot fully reveal the relationship and diversity of *P. effusa* isolates (Bourret *et al*., 2018). To further investigate the relationship between *P. effusa* isolates, we performed in-depth comparisons of mitochondrial and nuclear genome variation of the 26 *P. effusa* isolates used in this study. Using Illumina short-read data, we performed variant calling with GATK against the public nuclear genome of *Pe1* (Klein *et al*., 2020). In total, we identified 314,276 multiallelic variant sites that can be separated in 280,750 SNPs and 35,677 indels. On average, 35.8% of the variants per isolate are homozygous and 64.2% are heterozygous. Apart from *Pe1*, on average 1.53% of the variants per isolate are unique, 1.75% are shared by only two, and 19.67% are shared between all isolates (Supplementary Figure 5; Supplementary Table 1A). To uncover the evolutionary relationship of the 26 *P. effusa* isolates, we used 260,616 biallelic SNPs to determine the pairwise nuclear genome nucleotide diversity for each isolate, as a percentage of difference over the entire genome. The average nucleotide diversity (nd) between all isolates was 0.083% (0.008% standard deviation), with the closest related isolates being *US-11* and *US-13b* (0.055% nd), and the most distant isolates being *Pe6* and *Pe14* (0.097% nd). Based on these observations, we assigned three genomic clusters of closely related isolates that have been denominated to different races: Cluster i contains *Pe1, Pe2*, and *Pe3* (0.056% average nd); Cluster ii contains *Pe4* and *Pe6* (0.071% nd); and Cluster iii contains *Pe12, Pe14, R14, US-11*, and *US-13b* (0.058% average nd). We observed high similarity between additional isolates, for example *R13* and *Pe13* (0.066% nd), which were not assigned into separate clusters because they are designated to the same race (Figure 4A).

**Figure 4.**
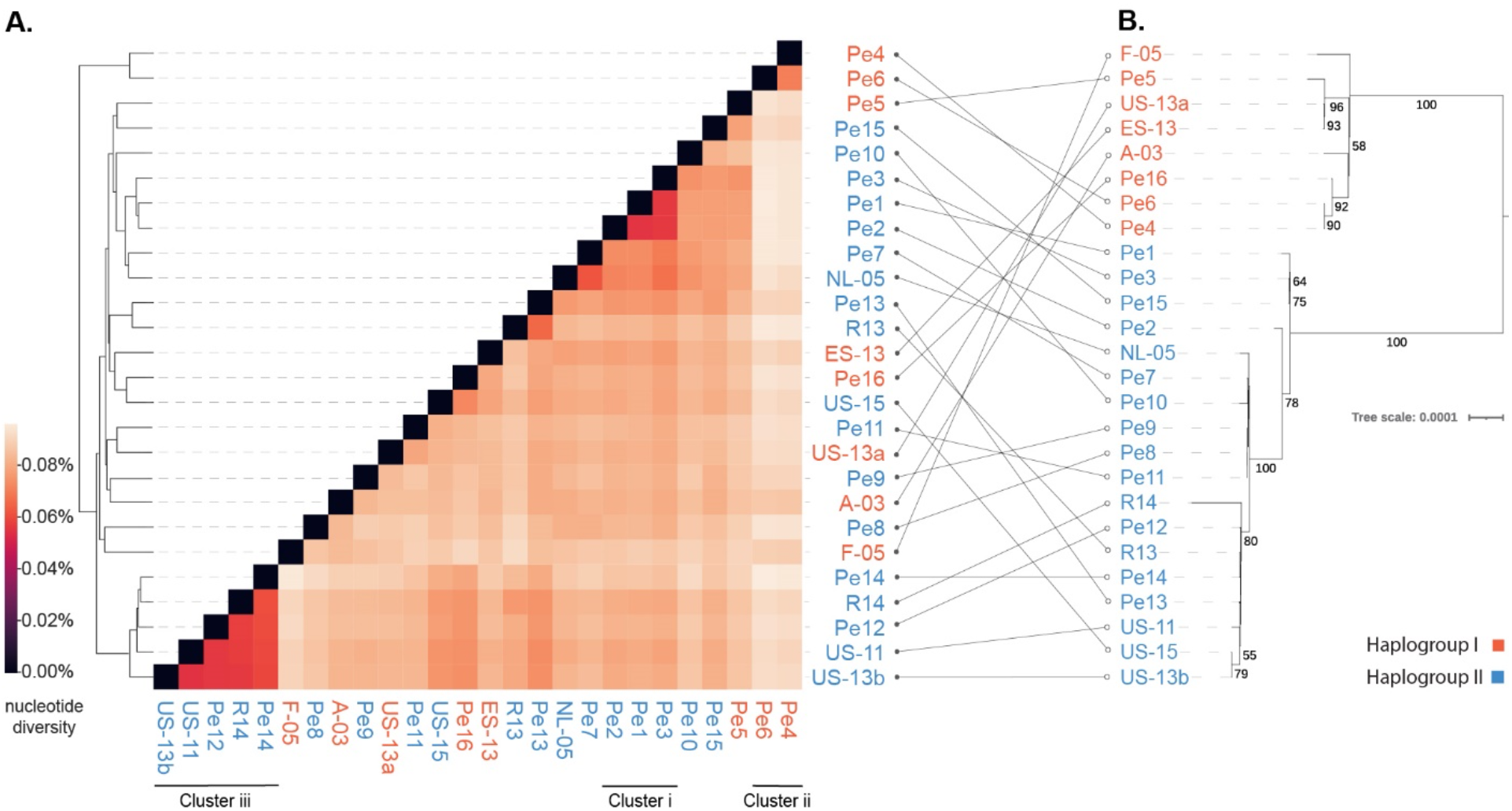
Nuclear and mitochondrial variation discordance in *Peronospora effusa (Pe)*. **A)** Hierarchically clustered heatmap of pairwise nuclear genome nucleotide diversity. Three distinct clusters formed by related isolates belonging to different races are highlighted. **B)** From the right-to the left, maximum-likelihood phylogeny based on the mitochondrial genome sequences of the 26 *P. effusa* isolates, rooted on the beet downy mildew (*Peronospora farinosa* f. sp. *betae, ES-15*). Concordance between this phylogeny and the hierarchical clustering shown in A) is highlighted by connecting lines between isolates.

To further explore the relationships between the isolates, we directly compared their similarity based on the nuclear variation with that identified from the mitochondrial phylogeny (Figure 4B). We observed that the ten isolates that cluster based on genome-wide nuclear similarity (>0.073% nd) also clearly cluster based on the mitochondrial phylogeny and belong to the same mitochondrial haplogroup (Figure 4B). For example, all isolates assigned to Cluster i (*Pe1, Pe2*, and *Pe3*) belong to Haplogroup II, while all the isolates assigned to Cluster ii (*Pe4* and *Pe6*) belong to Haplogroup I. By contrast, we observed clear discordance between nuclear clustering and the mitochondrial phylogeny for the 16 remaining isolates (Figure 4B). For example, isolates with high genome-wide nucleotide similarity such as *US-15* and *Pe16* (0.073% nd), *Pe5* and *Pe15* (0.078% nd), or *Pe11* and *US-13a* (0.079% nd) belong to different mitochondrial haplogroups.

### Mitochondrial and nuclear variation suggests that sexual recombination contributes to the evolution of resistance-breaking *Peronospora effusa* isolates

To explain the discordance between nuclear clustering and mitochondrial phylogeny of *P. effusa*, we further explored the genetic relatedness between *P. effusa* isolates. Based on the previously generated diversity matrix (260,616 biallelic SNPs), we explored the genetic differences by principal component analysis (PCA); the first principal component explains 14.14% of the observed variation and the second 13.22%, while the third principal component only explains 7.01%. When visualizing the first two principal components, we observed the same three distinct clusters that were previously recovered based on the genome-wide nuclear diversity (Figure 5A). The remaining isolates are localized in the middle of the PCA, sharing genetic material but did not clearly cluster in a distinct group.

**Figure 5.**
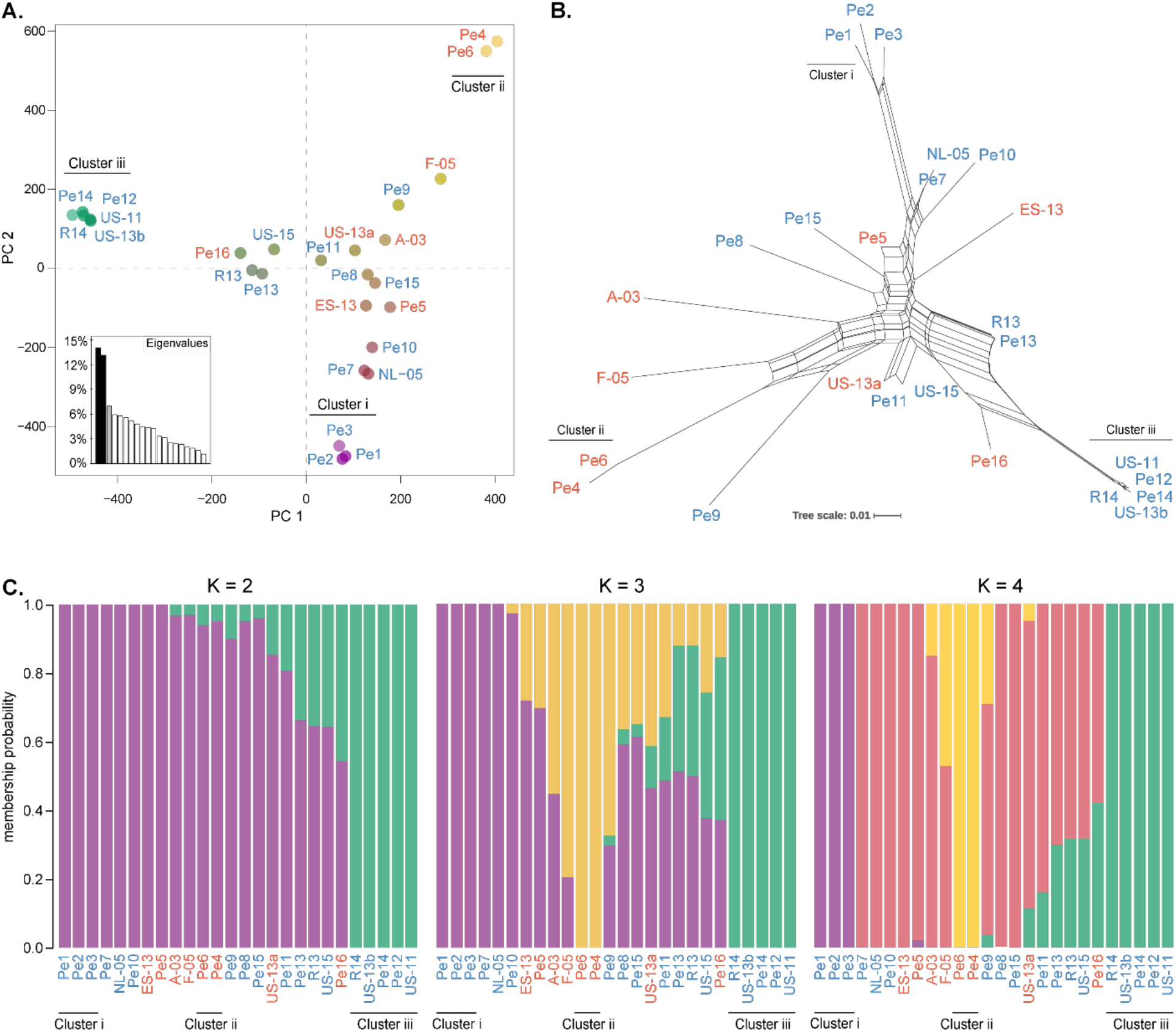
*Peronospora effusa* evolves by both asexual propagation and sexual recombination. Visualisations are constructed based on a distance matrix of genome-wide nucleotide differences, containing 260,616 biallelic sites for A and B and 169,016, minor allele frequency-filtered biallelic sites for C. The isolates are coloured based on their mitochondrial haplogroup (Haplogroup I in orange and Haplogroup II in blue), and the previously identified three distinct clusters of closely related isolates belonging to different races are highlighted and coloured similarly between A and C (Figure 4A). **A)** Principal component analysis; the axes depict the first two principal components that collectively explain 27.36% of the observed variation. **B)** Neighbor-net phylogenetic network; the branch lengths are proportional to the calculated number of substitutions per site. The parallel edges connecting different isolates indicate conflicting phylogenetic signals. **C)** STRUCTURE analysis for two, three, and four genetic sub-populations (K), without any prior assumptions for the grouping of isolates. Vertical bars represent *P. effusa* isolates, and the proportion of the coloured segments indicates the probability for the isolate to belong to *K* genetic sub-populations.

To better visualise the complex phylogenetic relationship of *P. effusa* isolates, we performed neighbor-net phylogenetic network analyses (260,616 biallelic SNPs), which is well-suited to illustrate the complex evolutionary relationships among different isolates in the presence of phylogenetic conflicts caused by, for instance, sexual recombination, hybridization, or horizontal gene transfer (Galtier *et al*., 2009). The neighbor-net phylogenetic network shows clear evidence of phylogenetic conflicts (Figure 5B), for instance between *Pe13* and *Pe5* that localized close to the centre of the network. *P. effusa* is known to reproduce sexually (Kandel *et al*., 2020), and thus, the most likely explanation for phylogenetic conflicts observed in our data is sexual recombination, which is further supported by the PHI test for recombination (260,616 biallelic SNPs, P < 0.0001) (Bruen *et al*., 2006). In contrast, the above defined three *P. effusa* nuclear clusters are connected to the main network with relatively long branches, suggesting that they accumulated a substantial number of cluster-specific variants (e.g., *Pe4* and *Pe6* uniquely share 5,029 (3.5%) variants), and thus, at least for some time, these isolates likely evolved without recombination with isolates outside of the respective cluster.

To further examine the exchange of genetic material between these isolates, we analysed the variation of *P. effusa* isolates using STRUCTURE, a population structure inference approach that uses Bayesian clustering based on allele frequencies to assign isolates to a pre-defined number of populations (K) (Linck and Battey, 2019). We performed STRUCTURE analysis with two, three, and four predefined sub-populations (K=2, K=3, K=4) (Figure 5C); any analysis with K>4 had identical results with K=4. When assuming two sub-populations (K=2), isolates belonging to both Cluster i and Cluster ii, and belonging to both haplogroups are present in a single sub-population (purple), thus their variation is not fully explained. When the isolates are divided into three and four sub-populations (K=3, K=4), the same three distinct clusters (i, ii, iii) that were previously identified by phylogenetic analysis were also assigned into three distinct sub-populations (purple, yellow, green) (Figure 4 and 5). This division in these three distinct sub-populations further confirms that the isolates in each cluster did likely reproduce asexually. In the case of K=3, all isolates belonging to the mitochondrial Haplogroup I are either a part of nuclear Cluster ii (yellow) or a recombination of nuclear Cluster ii with other clusters. For K=4, isolates that previously shared genetic material are now assigned to a distinct sub-population (red), which contains isolates belonging to both mitochondrial haplogroups. The presence of both mitochondrial haplogroups suggests that the isolates assigned to the red sub-population but have different virulence spectrum, do not have a common origin, but have likely evolved after one or more sexual recombination events between Cluster i and ii isolates. For example, for K=3 we observed that *Pe5* contains genetic information that is partially shared with Cluster i and ii (purple and yellow). We similarly observed genetic combinations of Clusters i, ii, and iii for some of the isolates (Figure 5C). For example, for K=3, *Pe13* is represented as a mix of the three defined sub-populations. This pattern could have emerged from multiple sexual recombination events involving different isolates of all three defined genetic sub-populations. We also observed a large variation in the number of unique variants for each isolate (Supplementary Figure 5). For *F-05*, for example, we observed 3,630 (2.79%) unique variants, which suggests that this isolate likely evolved asexually after the possible sexual recombination event. In contrast, we only observed 910 (0.73%) unique variants in *Pe11*, which suggests more recent sexual recombination events.

## Discussion

In the last two decades, new races of the spinach downy mildew *Peronospora effusa* (*Pe*) have been rapidly emerging, and thus far 19 *P. effusa* races have been denominated; the two latest ones have been added in 2021 (Correll and Smilde, 2021). However, we know little about the processes contributing to the diversity and evolution of different *P. effusa* races. To better understand the molecular differences between *P. effusa* races, we here described the mitochondrial and nuclear relationships between 16 denominated races and eight pathotypes of *P. effusa*; for the most recent races (*Pe* 17-19) no genomic data was yet available. Based on mitochondrial data, *P. effusa* isolates can be divided into two clear groups based on the sequence and structure. Based on nuclear data, we observed that a subset of isolates could be described by three distinct groups, which are in concordance with their mitochondrial phylogeny, in turn, suggesting that these isolates likely evolved through asexual reproduction. The remaining 16 isolates display discordance between nuclear and mitochondrial clustering as well as signs of shared genetic material, which uncovers that also sexual reproduction contributed to the emergence of *P. effusa* races.

Newly found *P. effusa* isolates are assigned to races based on their ability to infect a defined set of differential spinach lines (Supplementary Figure 1). The phenotyping and maintenance of *P. effusa* races, and the development of differential spinach lines is handled by the International Seed Federation (Correll *et al*., 2015) (Supplementary Table 4). The 24 isolates sequenced in this study have been carefully maintained and screened based on their phenotypes since their date of isolation, to avoid mixtures of multiple races (Feng et al., 2014). As it has been shown in other obligate biotrophic plant pathogens, what is described as an isolate could be a population of genetically distinct isolates with identical phenotypes, rather than a single genetically distinct isolate (Barsoum *et al*., 2020). However, based on our genome sequencing data, we are unable to find any indication of mixed genotypes in a single isolate. Based on our variant calling analysis against the mitochondrial and nuclear genome of *Pe1*, we did not observe any multiallelic sites for individual isolates, i.e., more than one allele for mitochondria and more than two alleles for nuclear genomes. In the same analysis of nuclear genome variants, we did not observe a significant deviation in allele frequency from the expected 50:50 of a diploid organism. Similarly, comparisons of isolates with the same phenotype (*Pe13* with *R13*, and *Pe14* with *R14*) shows that they have almost identical genotypes (0.066% and 0.061% nucleotide diversity, respectively), despite being isolated and sequenced independently (Fletcher *et al*., 2018). Thus, we conclude that our data does not suggest that any of our isolates is in fact a mixture of different genotypes.

Inverted repeats are known to form cruciform DNA structures that can cause chromosomal instabilities, leading to the formation of structural rearrangements (Achaz *et al*., 2003; Voineagu *et al*., 2008). Here, we observed inverted repeats to be associated with mitochondrial genome rearrangements in *P. effusa* as well as throughout related Peronosporaceae. Inverted repeats are genomic regions that are particularly challenging to assemble correctly when sequencing reads are not long enough to span these repeats (Wang *et al*., 2018). Inverted repeats in the *P. effusa* mitochondrial genome are 832 bp in size, making it impossible for short-read data to faithfully resolve mitochondrial chromosome structures. The *R13* and *R14* mitochondrial genomes display the structure of Haplogroup I isolates, even though these isolates are clearly assigned to Haplogroup II based on mitochondrial sequence variation. This discrepancy could be due to the short-read sequencing data used to reconstruct the *R13* and *R14* mitochondrial sequences (Fletcher *et al*., 2018). By contrast, we utilised long-read sequencing data to assemble mitochondrial genomes. The long reads span the inverted repeats, as corroborated by uniform read coverage throughout the mitochondrial sequence, which suggests that our mitochondrial assemblies of *Pe1, Pe4, Pe11*, and *Pe16* are accurate. The structural rearrangement that was initially observed in *Pe1* separates the two mitochondrial haplogroups of *P. effusa*. This rearrangement is not present in any queried oomycete, suggesting it is of relatively recent origin. Similarly, we can assume that the unique structural rearrangement in *Plasmopara viticola* is also formed by a relatively recent event. The DNA of these two recent rearrangements are flanked by inverted repeats, thus suggesting that genome instability caused by recombination of inverted repeats can drive mitochondrial diversity in Peronosporaceae. Because of this instability, we can foresee that rearrangements are likely to reoccur in this region. Thus, the inverted repeats itself would be a poor marker to robustly separate the defined haplogroups. To distinguish the mitochondrial haplogroups the inverted repeats could be augmented by the identified haplotype-specific, single-nucleotide variants.

The mitochondrial genes *cox2* and *nad1* have been previously used to separate *P. effusa* isolates into two district haplogroups that were found to be associated with geographical origin of the isolates tested (Choi *et al*., 2011). Here, we used 24 *P. effusa* isolates and robustly recovered these two distinct haplogroups based on *cox2* and *nad1*, whole-mitochondrial genome alignment, and structural variation. Even though most *P. effusa* isolates that were isolated outside the USA can be found in Haplogroup I, we were unable to establish a clear correlation between the mitochondrial haplogroup and the geographical origin of the isolates. Isolates with similar virulence spectrum are often simultaneously found at different geographical locations (Irish *et al*., 2003; Satou *et al*., 2006), which is likely due to the globalized spinach market where new spinach varieties are distributed and sold worldwide. Possibly, the sexually produced oospores of *P. effusa* that are abundant in spinach fields drive the repeated emergence of new races (Dhillon *et al*., 2020), followed by a rapid spread over a large area. Therefore, it is difficult to determine where a novel resistance-breaking isolate first occurred, and in our case, to know the exact geographical origin of the denominated *P. effusa* races.

*P. effusa* can reproduce both sexually and asexually (Kandel *et al*., 2020). It has been suggested that sexual reproduction is the main mechanism behind increased adaptability, while asexual reproduction allows the rapid propagation of successful genotypes thereby causing major disease epidemics (Drenth *et al*., 2019). As a result, sampling in the field during an epidemic will be greatly biased towards isolates that have been asexually reproducing, obscuring the role of sexual reproduction. Consequently, most *P. effusa* field isolates at a certain time and geographic location are the result of asexual reproduction (Lyon *et al*., 2016). Asexual reproduction can introduce significant phenotypic changes in resistant breaking *P. effusa* isolates (e.g., *Pe4* and *Pe6, Pe12 and Pe14*). Various mechanisms for asexual evolution have been documented in other oomycete and fungal plant pathogens. In *Phytophthora infestans* asexually evolving populations increase their genotypic variation by increase in ploidy and in gene copy number (Cooke *et al*., 2012; Knaus *et al*., 2020). Additionally, phenotypic differences can result from changes in gene expression (Cooke *et al*., 2012; Depotter *et al*., 2021). The forest pathogen *Phytophthora ramorum* has only been observed to reproduce asexually, with extensive mitotic recombination and gene copy number variation driving the genotypic diversity (Dale *et al*., 2019). In fungi it has been suggested that high levels of genotypic diversity can be generated by asexual reproduction through large structural variation (Seidl and Thomma, 2014; McDonald and Stukenbrock, 2016). For example, different races of the asexual fungal plant pathogen *Verticillium dahliae* evolve by extensive chromosomal rearrangements including large-scale gene losses (De Jonge *et al*., 2012; Faino *et al*., 2016; Chavarro-Carrero *et al*., 2021). In *P. effusa* loss of heterozygosity is suggested to contribute to the observed variation in asexually reproducing isolates (Lyon *et al*., 2016). From the available 13 *P. effusa* races that were discovered after 1990, nine have evolved through sexual recombination (*Pe5, Pe7* - *Pe11, Pe13, Pe15*, and *Pe16*). Thus, we hypothesise that in the period when *P. effusa* is under strong evolutionary pressure from the introduction of resistant spinach varieties, sexual reproduction has driven the fast emergence of new *P. effusa* races. Evolving sexually is often connected to a more stable genome structure. For example, in *Phytophthora infestans* sexually evolving populations are predominantly diploid and have less gene copy number variation than the asexual populations (Cooke *et al*., 2012). These findings show that there are multiple mechanisms for evolutionary adaptation, both with asexual and sexual reproduction (Seidl and Thomma, 2014).

Up to today, the genetic underpinnings of the race structure and evolution in *P. effusa* remain unknown. Based on the findings in other plant pathogenic oomycetes and fungi, it can be anticipated that the emergence of novel races is linked to genotypic changes in specific effector genes. Effector proteins are secreted by pathogens to modulate host physiology, often by deregulating immune responses, and establish infection. Resistant hosts produce immune receptors that can detect these effectors and trigger strong immune responses thereby stopping colonization. Effector genes often localize in repeat-rich genomic regions that are notoriously challenging to assemble with short-read sequencing that has been commonly used until recently (Gibriel *et al*., 2016). High quality, complete genome assemblies of multiple *P. effusa* isolates are thus invaluable to fully uncover the molecular underpinning of race emergence in *P. effusa* (Thomma *et al*., 2016). The newly generated high-quality genome assembly of *P. effusa* is much longer and more repetitive compared with previous genome assemblies (Fletcher *et al*., 2021), providing a valuable resource to start comparative research between races. To link genotypic variation to differences in resistance-breaking on spinach cultivars, emphasis should be given to the systematic mining of effector genes and their variation between genetically and phenotypically distinct *P. effusa* isolates. This research will greatly benefit from the here developed phylogenetic framework and insights in the proposed sexual recombination between different *P. effusa* races, which will facilitate our understanding of *P. effusa*-spinach interaction and assist in sustainable production of spinach through knowledge-drive resistance breeding.

## Experimental Procedures

### *Peronospora effusa* infection on soil-grown spinach and spore isolation

We sowed spinach plants in potting soil (Primasta, NL) and kept them under long-day conditions (16-hour light, 21 °C). Two to three weeks after germination, the spinach plants were inoculated with *P. effusa* by spraying them with *P. effusa* spores suspended in water using a spray gun. Following inoculation, the lids of the plastic trays were sprayed with water and covered to keep the plants humid and dark. After 24 hours, we placed the plants under 9-hour light and 16 °C. The lids of the boxes were again sprayed with water 7-10 days after inoculation, creating a humid environment that promotes the sporulation of *P. effusa*.

To harvest *P. effusa* spores for Oxford nanopore sequencing, we collected leaves with sporulating *P. effusa* from spinach plants and placed them in a glass bottle with tap water. The spores were brought into suspension by shaking the bottle vigorously. Soil and other large contaminants were removed by filtering the spore suspension over a 50-μm nylon mesh filter (Merck Millipore, USA). To remove small biological contaminants, the remaining filtrate was first filtered over a 11-μm nylon mesh filter (Merck Millipore, USA) using the Merck™ All-Glass Filter Holder (47 mm) and a vacuum pump. Afterwards, the remaining spores were resuspended in the All-Glass Filter Holder using 100 mL of autoclaved tap water and filtered.

### High-molecular weight DNA extraction protocol

To isolate high-molecular weight (HMW) DNA, the collected *P. effusa* spores were freeze-dried and lysed using glass beads in a tissue lyser (2 × 30 seconds on 30 Hz). The lysed spores were incubated in extraction buffer (3% CTAB, 2% PVP, 1.25 M NaCl, 200 mM Tris-HCl, pH 8.5, 25 mM EDTA, pH 8.0) for 30 minutes at 65 °C, gently inverting the tube several times every 5 minutes. Then RNase was added, and the sample was incubated for 30 minutes at 37 °C, after which 0.5% 2-Mercaptoethanol was added, followed by another 15-minute incubation at 37 °C. HMW DNA was isolated from the lysate by subsequent phenol/chloroform/IAA extraction, chloroform/IAA washing, RNase treatment, another phenol/chloroform/IAA extraction, chloroform washing, and lastly, isopropanol precipitation.

DNA purity was determined with Nanodrop (Thermo Fisher Scientific, USA) to measure A260/280 and A260/230 ratios. DNA concentrations were estimated using a fluorimeter and fluorescent DNA-binding dye (Qubit™ dsDNA BR Assay Kit; Thermo Fisher Scientific, USA), according to the manufacturer’s protocol. DNA integrity was confirmed by 0.4% agarose gel electrophoresis performed at 4 °C at 20 V overnight.

### Genome sequencing using Oxford Nanopore

We obtained long-read sequencing data for four *P. effusa* isolates (Pe1, Pe4, Pe11, Pe16) with Oxford Nanopore sequencing technology (Oxford Nanopore, UK). The ligation-based sequencing kit from Oxford Nanopore was used for library preparation (ONT - SQK-LSK109; Oxford Nanopore, UK) following the manufacturer’s protocol, apart from DNA-nick repair. This was performed as described by the protocol of Illumina sequence library preparation (kit: NEB - M6630; protocol: E6040; Illumina, USA). We used a Nanopore MinION flowcell (R10) for real-time sequencing, and base-calling of the raw long-read sequencing data was performed using Guppy (version 4.4.2; default settings). The raw long-read sequencing data were checked for contamination using Kraken2 (version 2.0.9; default settings).

### Genome sequencing using Illumina

We obtained paired-end short-read sequencing data of 24 *P. effusa* isolates and the *Peronospora farinosa* f. sp. *betae* isolate *ES-15* on the Illumina sequencing platform (Illumina, USA). Sequencing libraries were constructed with the Illumina TruSeq DNA PCR-Free kit. Fragment-size distribution was determined before and after the library preparation using the Agilent Bioanalyzer 2100 with HS-DNA chip (Agilent Technologies). The final library size was approximately 550 bp). Libraries were paired-end sequenced (two times 150-bp reads) on the Illumina NextSeq platform (Utrecht Sequencing Facility) in high output mode.

### Mitochondrial genome assembly and annotation

We selected on average 155 Mb of long-read sequences from the four sequenced *P. effusa* isolates based on their sequence alignment to 26 publicly available oomycete mitochondrial genomes. To this end, we aligned the long-read sequences to these 26 mitochondrial genomes using mimimap2 (version 2.17) (Li, 2018) with recommended settings for Nanopore reads. The extracted reads were subsequently assembled with Canu (version 2.0) (Koren *et al*., 2017) with recommended settings for the assembly of Nanopore reads. We mapped the high-quality Illumina paired-end short-reads of the four *P. effusa* isolates to their respective draft assembly with BWA-mem (version 0.7.17; default settings) (Li and Durbin, 2009), and sequencing errors in the draft mitochondrial genome assembly were corrected with Pilon (version 1.23; default settings) (Walker *et al*., 2014). Genes were annotated according to the universal genetic code and based on sequence similarity with known mitochondrial genes using GeSeq (Tillich *et al*., 2017). Inverted repeats were detected by self-alignment of mitochondrial genome sequences and visualised using Circoletto (version 07.09.16) (Darzentas, 2010).

Clinker (version 0.0.12) (Gilchrist and Chooi, 2021) was used to visualise the mitochondrial genome structure, by aligning the mitochondrial protein sequences of *Pe1* together with 17 Peronosporaceae species and using those alignments as ankers to visualise the order of genes in the genomes.

### Variant calling on mitochondrial and nuclear genomes

We used Illumina paired-end genomic short-read data for 26 *P. effusa* isolates to discover single-nucleotide variants. We first filtered short reads by quality using Trimmomatic (version 0.39-1; MINLEN:36 LEADING:3 TRAILING:3 SLIDINGWINDOW:4:15) (Bolger *et al*., 2014) and removed Illumina sequencing artefacts with Fastp (version 0.20.0) (Chen *et al*., 2018). The filtered short-read data were aligned to the newly generated mitochondrial genome sequences or to the publicly available nuclear genome assemblies of *Pe1* with BWA-mem (version 0.7.17; default settings) (Li and Durbin, 2009; Klein *et al*., 2020). We identified single-nucleotide variants (SNPs) using the GATK joint variant calling pipeline (version 4.1.9.0) following the best practices for germline short-variant discovery. We filtered individual variants based on the hard-filtering best practices of GATK: Quality by depth < 4.0, Fisher Strand > 60.0, Strand Odds Ratio > 3.0, RMS Mapping Quality < 20.0, Mapping Quality Rank Sum Test < -3.0, Read Position Rank Sum Test < -1.0, Read Position Rank Sum Test > 3.5.

### Mitochondrial genome phylogeny

The SNPs identified in each of the 26 *P. effusa* isolates were incorporated in the *Pe1* mitochondrial genome sequence to reconstruct an isolate-specific mitochondrial sequence with FastaAlternateReferenceMaker (GATK version 4.1.9.0; default settings). The resulting isolate-specific sequences were aligned using MAFFT (version 7.471; default settings) (Nakamura *et al*., 2018) and a maximum-likelihood genome phylogeny was constructed using IQ-TREE (version 1.6.12; default settings) (Nguyen *et al*., 2015). Branch support was evaluated using 1,000 bootstrap replicates. Similarly, we reconstructed a maximum-likelihood phylogeny based on the sequences of the mitochondrial genes *cox2* and *nad1* that were extracted from the mitochondrial genomes of 18 Peronosporaceae.

### Nuclear genome nucleotide diversity

Based on the data for all 26 *P. effusa* isolates, we generated a joint VCF file with both variant and invariant sites, using the variant calling pipeline described above with the added option “-all-sites” in the tool GenotypeGVCFs (GATK version 4.1.9.0). We calculated the nucleotide diversity between the *P. effusa* isolates with pixy (version 1.0.4; --chunk_size 1000000) (Korunes and Samuk, 2021). We performed hierarchical clustering of the genome-wide nucleotide diversity and visualized the data as a hierarchically clustered heatmap with seaborn (version 0.11.1) (Waskom, 2021).

The single nucleotide variants of the 26 *P. effusa* isolates were transformed into a distance matrix with PGDSpider (version 2.1.1.5) (Lischer and Excoffier, 2012), which was used for the following analysis: first, we performed a principal component analysis and visualised the first two principal components on a scatterplot in RStudio (libraries: vcfR version 1.12.0, adegenet version 2.1.3) (Jombart and Ahmed, 2011; Knaus and Grünwald, 2017). Second, we constructed a decomposition network using the Neighbor-Net algorithm with SplitsTree (version 4.17.0) (Huson and Bryant, 2006). We calculated the branch confidence of the network using 1,000 bootstrap replicates. PHI recombination test was calculated using phipack (version 1.1; default settings) (Bruen *et al*., 2006). The single nucleotide nuclear variants of the 26 *P. effusa* isolates were additionally filtered for Minor Allele Frequency < 0.1 (169,016 biallelic SNPs), and transformed into the STRUCTURE format with PGDSpider (version 2.1.1.5). We performed STRUCTURE analysis using fastStructure (version 1.0; default settings) (Raj *et al*., 2014) with various k values. The results were visualised on a stacked bar-plot in RStudio (libraries: vcfR version 1.12.0, poppr version 2.9.0) (Kamvar *et al*., 2015; Knaus and Grünwald, 2017).

## Acknowledgments

This project was financially supported by the TopSector TKI Tuinbouw & Uitgangsmaterialen, the Netherlands, and four private breading companies: Enza Zaden Research & Development BV, Pop Vriend Research B.V., Rijk Zwaan Breeding, and Syngenta.

## Data availability

For this study we sequenced 16 denominated races of *P. effusa*. Cultures of these can be requested for research from Naktuinbouw, the Netherlands (Correll *et al*., 2015) (Supplementary Table 4). The raw sequence data and genome assemblies generated in this study are available at NCBI under BioProject PRJNA772192.

