## Supplementary Figures for "Sexual reproduction contributes to the evolution of resistance breaking isolates of the spinach pathogen *Peronospora effusa*"

### Supplementary Tables

(provided in separate excel files)

**Supplementary Table 1. Metrics of sequencing data used in this project. A.** Illumina paired-end whole genome sequencing of 26 *P. effusa* isolates and quality control (yellow); variant calling of nuclear genome based on the *Pe1* reference (blue); variant calling of mitochondrial genome based on *Pe1* reference (green). **B.** Nanopore whole genome sequencing of four *P. effusa* isolates (yellow); mitochondrial genome assemblies generated with the nanopore sequencing data (blue).

**Supplementary Table 2.** List of the species that their mitochondrial genome assembly was used to bait the reads for the mitochondrial genome assemblies of *Pe1*, *Pe4*, *Pe6*, and *Pe11*.

**Supplementary Table 3.** Gene annotation of *Pe1* mitochondrial genome generated with GeSeq.

**Supplementary table 4.** Details of the *P. effusa* isolates that have been sequenced in this study.

| | Pe: 1 | Pe: 2 | Pe: 3 | Pe: 4 | Pe: 5 | Pe: 6 | Pe: 7 | Pe: 8 | Pe: 9 | Pe: 10 | Pe: 11 | Pe: 12 | Pe: 13 | Pe: 14 | Pe: 15 | Pe: 16 |
|---|---|---|---|---|---|---|---|---|---|---|---|---|---|---|---|---|
| **Viroflay** | + | + | + | + | + | + | + | + | + | + | + | + | + | + | + | + |
| **NIL 5** | - | - | + | + | + | + | + | + | + | + | + | + | + | + | + | + |
| **NIL 3** | - | + | - | + | - | + | + | - | - | + | - | - | + | - | + | - |
| **NIL 4** | - | - | - | - | + | + | + | + | + | + | + | + | + | + | - | + |
| **NIL 6** | - | + | - | - | - | + | - | + | + | + | - | + | - | + | - | - |
| **NIL 1** | - | - | - | - | - | - | - | + | - | + | - | + | (-) | + | - | - |
| **NIL 2** | - | - | - | - | - | - | - | - | - | - | + | + | + | + | - | + |
| **Pigeon** | - | - | - | - | - | - | - | - | - | - | - | - | - | + | - | + |
| **Caladonia** | - | - | - | - | - | - | - | - | - | - | - | - | - | - | + | - |
| **Meerkat** | - | - | - | - | - | - | - | - | - | - | - | - | - | - | - | + |
| **Hydrus** | - | - | - | - | - | - | - | - | - | - | - | - | - | - | - | - |

**Supplementary Figure 1.** Breaking of resistance on spinach differential set from the 16 *Peronospora effusa* designated races.

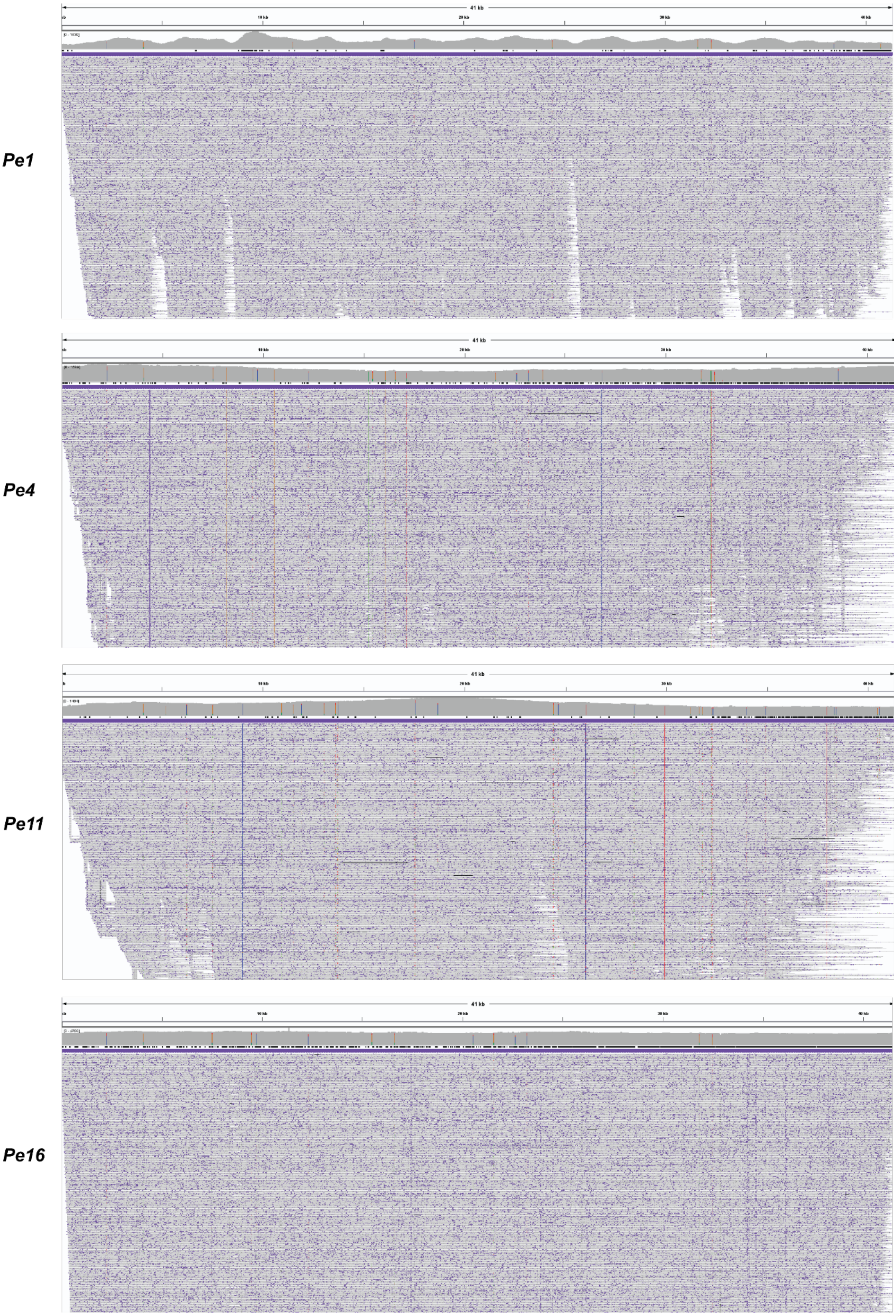


**Supplementary Figure 2. Coverage of the mitochondrial genome assemblies of *Peronospora effusa* races 1, 4, 11, and 16 (*Pe1*, *Pe4*, *Pe11*, *Pe16*).** For each race, the sequenced nanopore long-reads were mapped back to their mitochondrial genome assembly, filtered for mapping quality 20 and visualised in IGV (version 2.8.13).

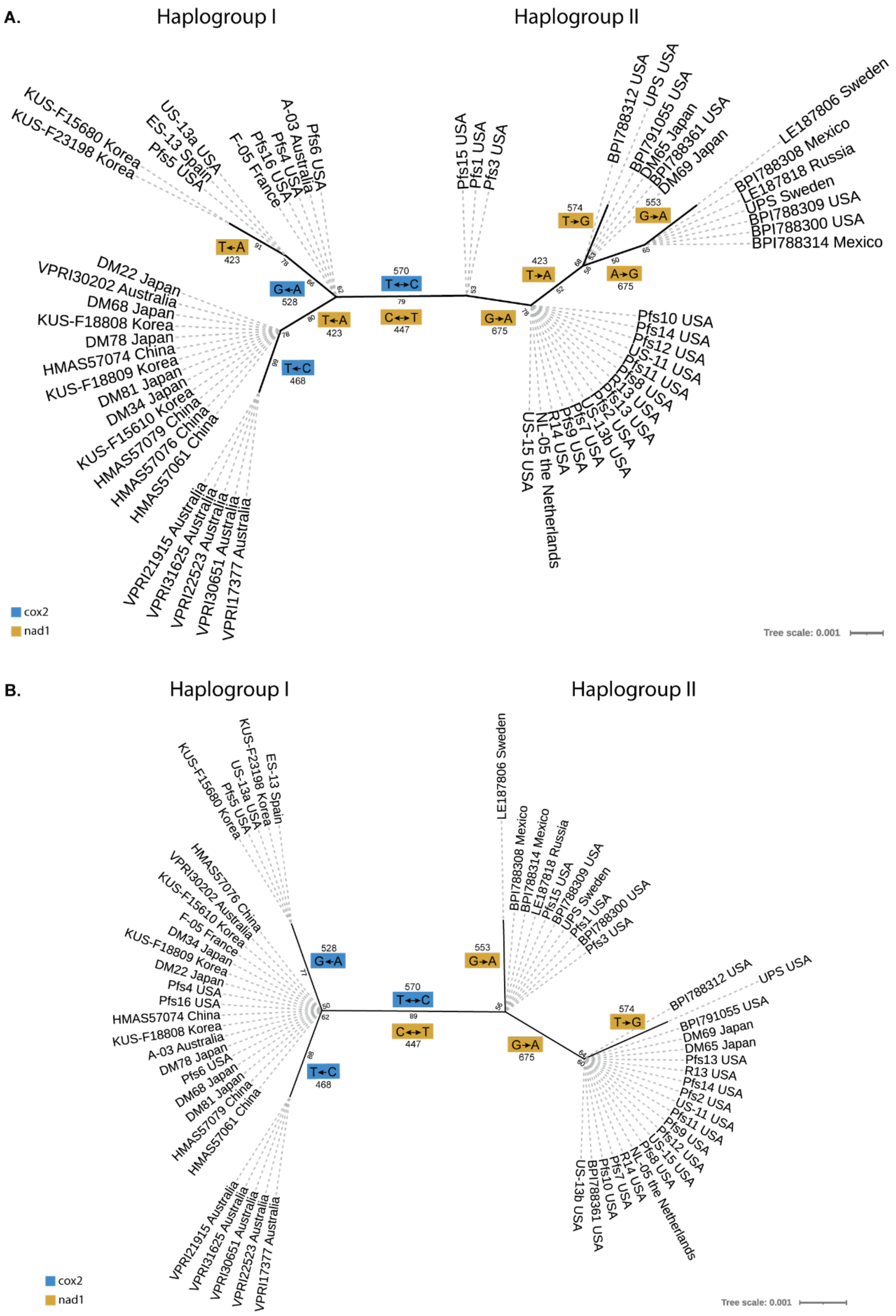


**Supplementary Figure 3.** ***Peronospora effusa*** **(here named** ***Pfs*****) phylogeny of the** ***cox2*** **and** ***nad1*** **mitochondrial genes reveals two distinct groups.** Unrooted maximum-likelihood phylogeny based on a 500bp sequence of the *cox2* and *nad1* genes from our 24 isolates together with the previously analysed 33 isolates from the global population (Choi et al., 2011) as well as *R13* and *R14* (Fletcher et al. 2018). The specific substitutions and their location separating each node in the trees are depicted in coloured boxes, blue for *cox2* and orange for *nad1*. **A)** A substitution (T to A) in the third nucleotide of the *nad1* subsequence (position 423 in the whole sequence) separates the 26 isolates sequenced with Illumina technology and the 33 isolates sequenced with SANGER technology (Choi et al., 2011), suggesting it's due to a sequencing error. **B)** The first 3 nucleotides of the *nad1* subsequence was excluded because of the possible sequencing error.

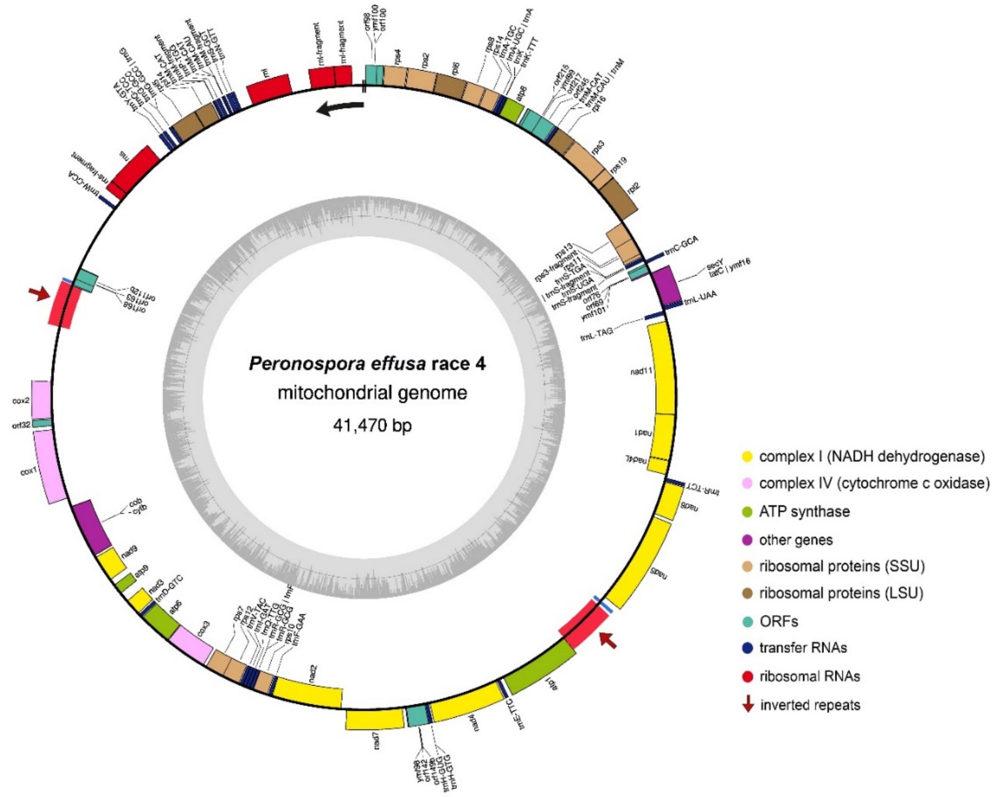


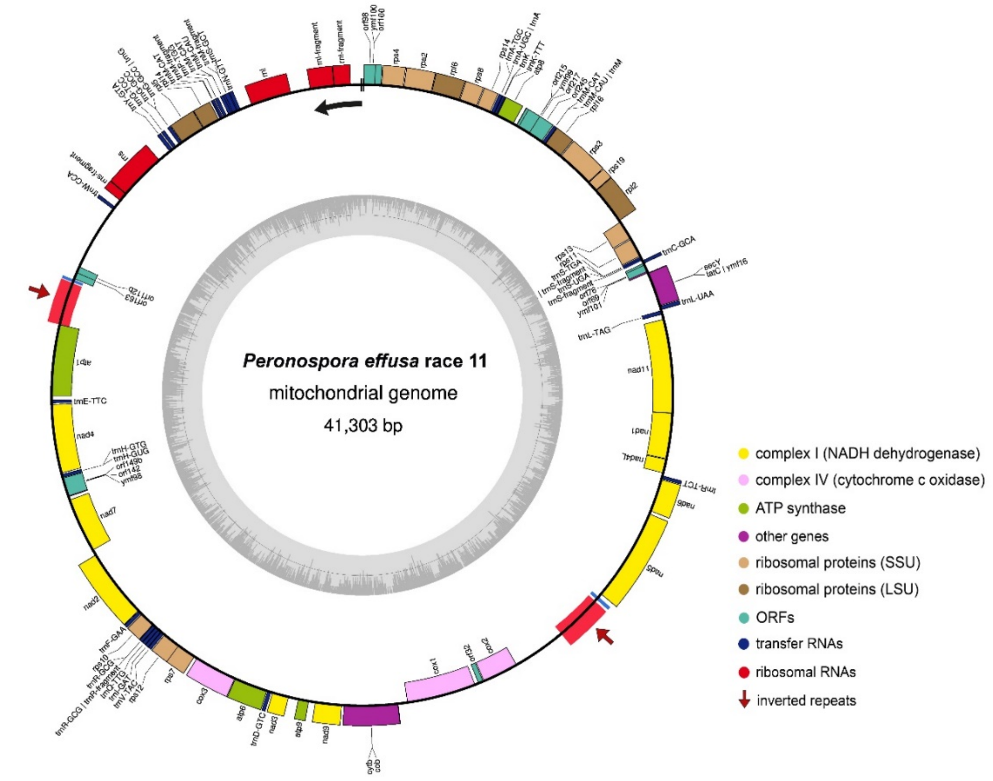

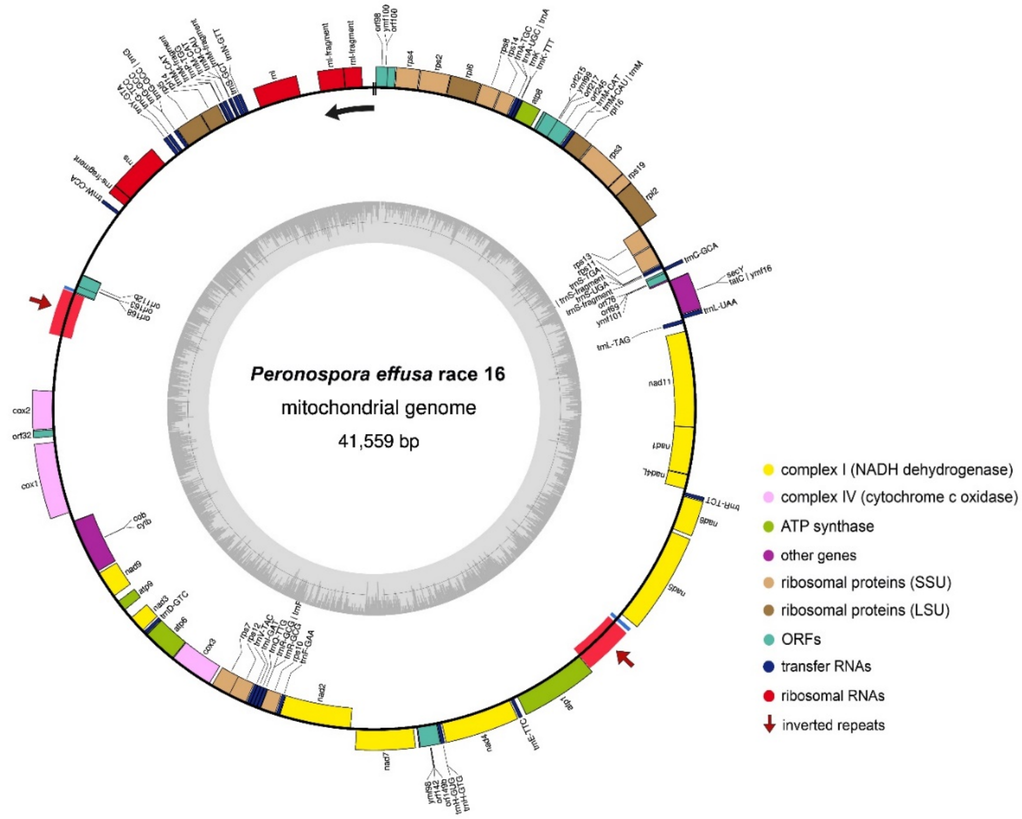


**Supplementary Figure 4. Mitochondrial genome annotation of *Peronospora effusa* races 4, 11, and 16 (*Pe4*, *Pe11*, *Pe16*).** Protein-coding genes, tRNA, rRNA, and other open-reading frames (ORFs) are shown along the outer ring (positive strand is the outside of the ring and the negative strand is the inside). The regions of inverted repeats are highlighted in red with arrows and are present in both strands. The inner ring depicts the GC content. The start and end of the linear representations (Figure 2B, Figure 3B) of the circular genome assembly is indicated with two black lines, with the arrow indicating the direction.

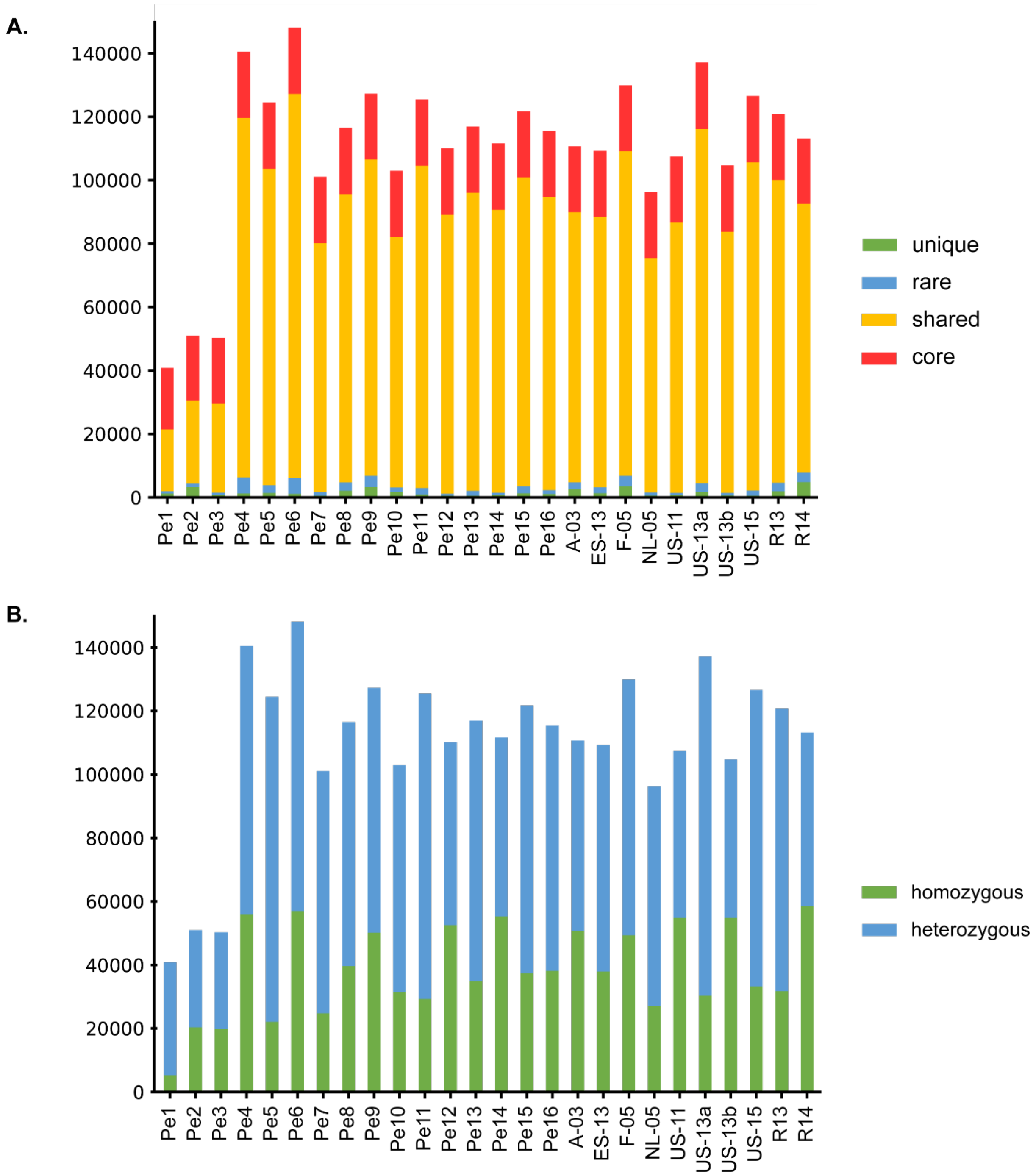


**Supplementary Figure 5. Nuclear genome variation of *Peronospora effusa* isolates.** Bar plots depicting the number of short variants (SNPs and single nucleotide INDELs) for each isolate in comparison to the *Pe1* reference genome. **A)** Distinction of variants in: unique, shared by only two (rare), shared by more than half of the isolates (shared), and shared between all isolates other than *Pe1* (core). **B)** Distinction of variants in homozygous and heterozygous.
